# Repeated extrinsic rewards following retrieval practice facilitate later memory

**DOI:** 10.1101/2025.07.01.662557

**Authors:** Devyn E. Smith, Amanda M. Smith, Hannah R. Buras, Nicole M. Long

## Abstract

The anticipation of extrinsic reward facilitates memory formation. However, it is unclear how reward following memory retrieval influences the information that is retrieved and later remembered. Here, we conducted four behavioral experiments (N=42 male/female young adults per experiment) in which we manipulated retrieval practice reward delivery. Across all experiments, participants studied word-image pairs and then completed two rounds of retrieval practice, followed by a final recognition test. Participants made vividness judgments during retrieval practice and in three of four experiments each response had a 50% chance of yielding positive feedback. We find that repeated rewards following retrieval practice facilitate later memory whereas low vivid retrieval practice impairs later memory. Together, these results suggest that the benefit of both retrieval practice and reward may be dependent on the strength of the memory that is retrieved.

## Introduction

Episodic memories may be reinforceable through extrinsic rewards (e.g. monetary compensation) as information that is valuable or rewarding is prioritized over information that is less rewarding (Loftus & Wickens, 1970; Dickerson & Adcock, 2018). The potential for future rewards enhances memory formation or encoding (Adcock, Thangavel, Whitfield-Gabrieli, Knutson, & Gabrieli, 2006). However, the extent to which reward *following* memory retrieval impacts the contents of retrieval and subsequent behavior is unknown. Prior behavioral work that has investigated test phase extrinsic reward has found mixed results (Shigemune, Tsukiura, Nouchi, Kambara, & Kawashima, 2017; Castanheira, Lalla, Ocampo, Otto, & Sheldon, 2022). However, these studies used anticipatory methods, which, like studies employing value directed remembering (Castel, Benjamin, Craik, & Watkins, 2002; Castel, 2007), have the potential to modulate the strategies that individuals use, rather than modulating the contents of retrieval. Thus, it is an open question how receiving a non-anticipated reward immediately following memory retrieval impacts processing and subsequent behavior. How extrinsic test-phase rewards impact retrieval is important because memory reinforcement may be better accomplished through direct reward of what is retrieved, rather than through manipulation of potential future reward, and may more closely mirror intrinsic responses to retrieval success (Speer, Bhanji, & Delgado, 2014; Smith & Long, 2024). The aim of this study was to investigate how extrinsic reward following memory retrieval impacts subsequent memory.

It is well established that associating a study item with a potential reward – e.g. reward that will be received if the item is remembered at test – impacts the likelihood that an item is later remembered (Loftus & Wickens, 1970; Marini, Marzi, & Viggiano, 2011; Elliott, Blais, McClure, & Brewer, 2020), with higher potential rewards leading to better subsequent memory. Enhanced subsequent memory for study items assigned a potential reward is driven by correlated activity between reward regions (e.g. ventral tegmental area, striatum) and the hippocampus, in which reward signals up-regulate hippocampal memory encoding mechanisms (Adcock et al., 2006; Wolosin, Zeithamova, & Preston, 2012). Given that the hippocampus supports both memory encoding and memory retrieval (Eichenbaum, 2004; Diana, Yonelinas, & Ranganath, 2007; Long et al., 2017), a similar interaction between the reward system and hippocampus during retrieval may also serve to enhance memory performance.

Practicing retrieval without any rewards is well known to improve later memory (Roediger & Karpicke, 2006; Karpicke & Roediger, 2008; Karpicke, 2012). However, incomplete or partial retrieval has the potential to negatively impact later memory (Detre, Natarajan, Gershman, & Norman, 2013). According to the non-monotonic plasticity hypothesis, a memory representation is weakened when it is moderately or partially reactivated (Poppenk & Norman, 2014). Thus, how vividly a memory is reactivated during retrieval practice will impact its later memory (Kuhl, Johnson, & Chun, 2013; Lee, Samide, Richter, & Kuhl, 2018). Potentially, rewards administered following retrieval may serve to up-regulate attention to the reactivated representation, strengthening a weak representation that would otherwise be forgotten.

Extant studies suggest that test phase rewards both improve and have no effect on memory. Shigemune and colleagues (Shigemune et al., 2017) had participants study words in high or low difficulty tasks and then complete a memory test. Prior to each test trial, participants were given a cue indicating that correctly recognizing a study item would result in a high or low reward. Hit rates (rates of correctly recognizing study items) were higher in the high compared to low reward condition in the high difficulty task. However, another study with a similar design (Castanheira et al., 2022) found no effect of potential reward during test. Additionally, as both of these studies measured the influence of reward on immediate memory performance – that is, how the potential for reward on the current trial impacted memory performance on the current trial – they were not positioned to demonstrate whether extrinsic rewards have a lasting impact on memory representations. Thus, the role of extrinsic test phase rewards remains limited.

A limitation of existing studies – regardless of whether potential reward is manipulated during the study or test phase – is that reward delivery is always anticipatory. In these motivated memory studies, individuals are aware prior to encountering a study or test stimulus that there is the potential to receive a reward for remembering that stimulus. Such a design will impact how upcoming stimuli are processed, as evidenced by studies employing value directed remembering (Castel et al., 2002; Castel, 2007). Both young and older adults can selectively attend to high value information and apply elaborative encoding strategies to high value items (Knowlton & Castel, 2022) and knowing how to capitalize on the value structure alters use of memory encoding strategies (Filiz & Dobbins, 2024). Thus, the discrepancy in the existing findings (Shigemune et al., 2017; Castanheira et al., 2022) may be driven by differences in strategy adoption. In contrast, non-anticipated rewards following retrieval are unlikely to consistently modify anticipatory strategy engagement. Unlike anticipatory rewards, post-retrieval reward may lead to alterations in memory representations. Thus, it is important to identify how extrinsic reward immediately following memory retrieval impacts the information that is retrieved.

Understanding the impact of post-retrieval reward is important because rewards can be both facilitatory and deleterious. High value rewards can have negative impacts on memory (Chung & Federmeier, 2023). Retrieval practice can facilitate memory but also increase errors to similar novel stimuli if general information common to both study items and lures is strengthened (McDermott, 2006; Lee et al., 2018). Presenting rewards during retrieval practice may or may not serve to facilitate later memory. As rewards can extend across related items (Oyarzún, Packard, de Diego-Balaguer, & Fuentemilla, 2016) and to inferred associations (Wimmer & Shohamy, 2012), rewards may promote erroneous or false memories. To the extent that participants reactivate category-level information during retrieval practice (e.g. Lee et al., 2018), rewards could reinforce category or gist-like representations over item-specific or verbatim representations (Brainerd & Reyna, 2002), which could lead to an increase in false memories for categorically-related lures presented during test.

Our hypothesis is that extrinsic reward following retrieval will reinforce the information that is retrieved and modulate subsequent memory. To test our hypothesis, we conducted four behavioral experiments in which we manipulated test phase reward delivery. Across all experiments, participants studied word-image pairs and then completed two rounds of retrieval practice, followed by a final recognition test. During retrieval practice, participants were given a word cue and instructed to bring to mind the associated image. They rated the vividness of their memory for the image on a scale from one (least vivid) to four (most vivid). In experiments 1-3, every response had a 50% chance of receiving reward feedback. To the extent that reward reinforces the information that is retrieved, we should find increased hit rates for reward items compared to not rewarded items.

## Materials and Methods

### Participants

168 native English speakers from the University of Virginia community participated, with 42 participants enrolled in each experiment (E1: 22 female; age range = 18-22, mean age = 19.2 years; E2: 27 female; age range = 18-21, mean age = 18.8 years; E3: 26 female; age range = 18-21, mean age = 19 years; E4: 25 female; age range = 18-21, mean age = 19.1 years). All participants had normal or corrected-to-normal vision. Informed consent was obtained in accordance with University of Virginia Institutional Review Board for Social and Behavioral Research and participants received class credit for their participation. Our sample size was determined *a priori* based on pilot data (E2, N = 14) described in the pre-registration report of this study (osf.io/gebm4). A total of 30 participants were excluded from the final dataset. Two (one each from E2 and E4) were excluded due to having a d’ *>* 2.5 SDs of the mean across the four experiments. Twenty-eight participants (E1: 8; E2: 5; E3: 10; E4: 5) were excluded due to failing to respond to all interleaved odd/even trials (see below) during the study phase. Thus data are reported for the remaining 138 (E1: 34, E2: 36, E3: 32, E4: 36) participants. All raw, de-identified data and the associated experimental and analysis codes used in this study are available via the Open Science Foundation (osf.io/y5ac8).

### Recognition Task Experimental Design

We conducted four recognition memory experiments (E1, E2, E3, E4) each with three phases (Figure 1) and manipulated test phase reward delivery between participants. Stimuli consisted of 1602 words, drawn from the Toronto Noun Pool (Friendly, Franklin, Hoffman, & Rubin, 1982) and three categories of images: 490 common objects (e.g., banjo), drawn from an image database with multiple exemplars per object category (Konkle, Brady, Alvarez, & Oliva, 2010), 96 famous faces (e.g., Britney Spears) and 96 famous scenes (e.g., Taj Mahal; Lee et al., 2018). From this set, 192 words and 288 images were selected for each participant. The images consisted of an equal number (96) of objects, faces, and scenes. Of the 288 images, a subset of 192 were presented in Phase 1, with 64 images drawn from each visual category. Trial counts by condition are shown in Figure 2. Only one exemplar per object category appeared during Phase 1 (e.g. one banjo). Word-image associations were randomly generated for each participant and randomly assigned to condition (e.g. target or lure, see below).

**Figure 1.**
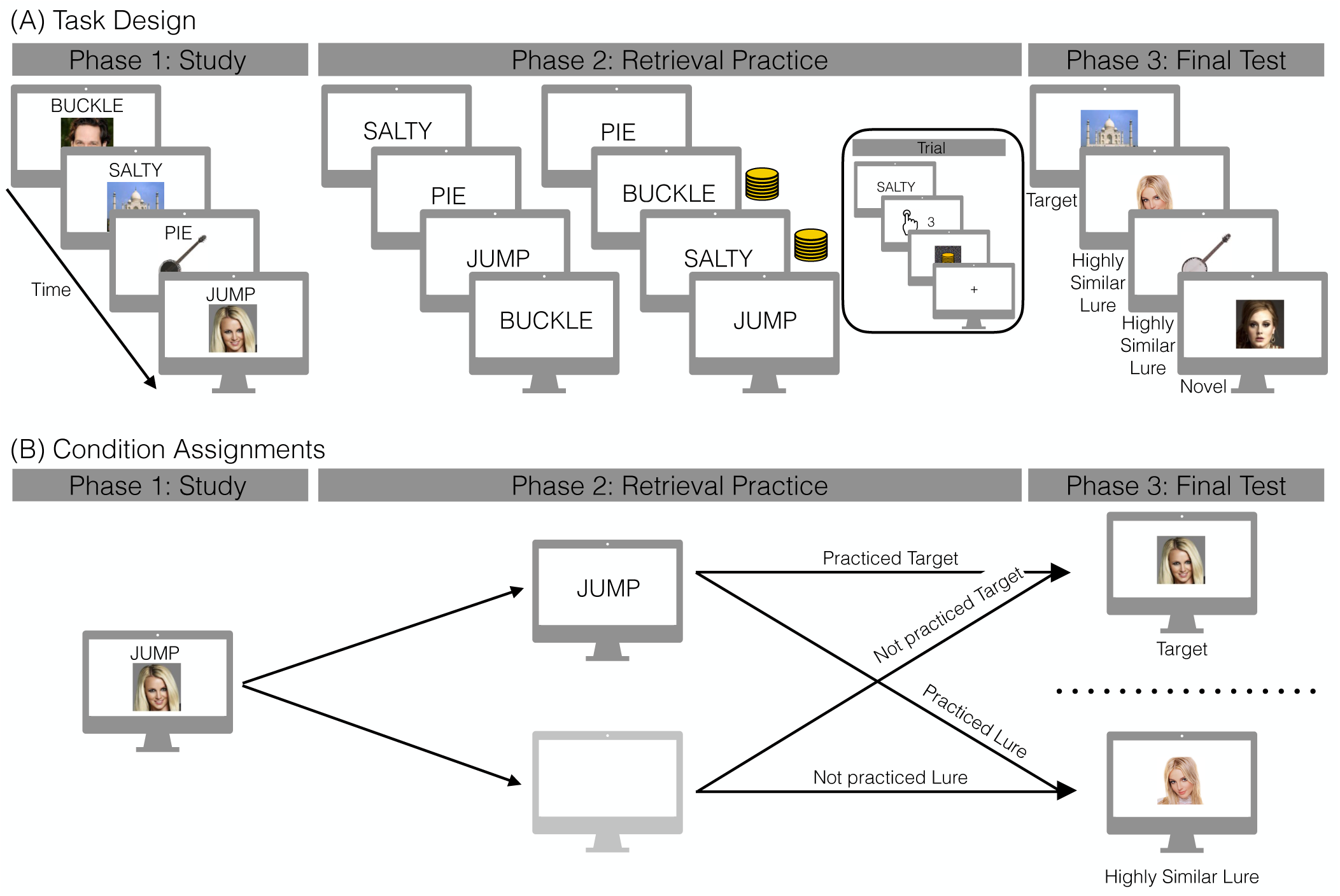
Task design and condition assignments. **(A)** During Phase 1, participants studied word-image pairs; images were from one of three categories: famous faces (e.g. Britney Spears), famous scenes (e.g. Taj Mahal), and common objects (e.g. banjo). In Phase 2, participants completed two rounds of retrieval practice. Participants saw an individual word and were instructed to bring to mind the image associated with each word and make vividness ratings on a scale from 1 to 4, with 1 being least vivid and 4 being most vivid. In E1, during the first round of retrieval practice, every response had a 50% chance of receiving reward feedback displayed as coins over a mask. The mask is shown alone on no reward trials. In E2, during the second round of retrieval practice, every response had a 50% chance of receiving reward. In E3, participants received rewards during both rounds of retrieval practice. In the first round, as in E1, every response had a 50% chance of receiving reward. If reward followed an item in the first round of retrieval practice, reward followed the same item in the second round. In E4, participants did not receive any reward during either round of retrieval practice. The temporal dynamics of a trial during retrieval practice are as follows: word is presented, participants rate the vividness of their memory for the image, then a reward could immediately follow. All participants then completed Phase 3, a final recognition memory test that included images only. Test probes included previously studied images (targets), highly similar lures (non-identical images depicting the same person, place, or object as the targets), and novel images. Participants made old or new judgements using a confidence rating scale from 1 to 4, with 1 being definitely new and 4 being definitely old. **(B)** We refer to both targets and lures as ‘practiced’ or ‘not practiced’ on the basis of whether the association (e.g. JUMP-Britney Spears was practiced or not (greyed monitor image) during Phase 2. Participants would only ever experience one condition per association, meaning that no participant would see both images of Britney Spears. These conditions can be further sub-divided on the basis of rewards and/or vividness rating during Phase 2 (see Figure 2).

**Figure 2.**
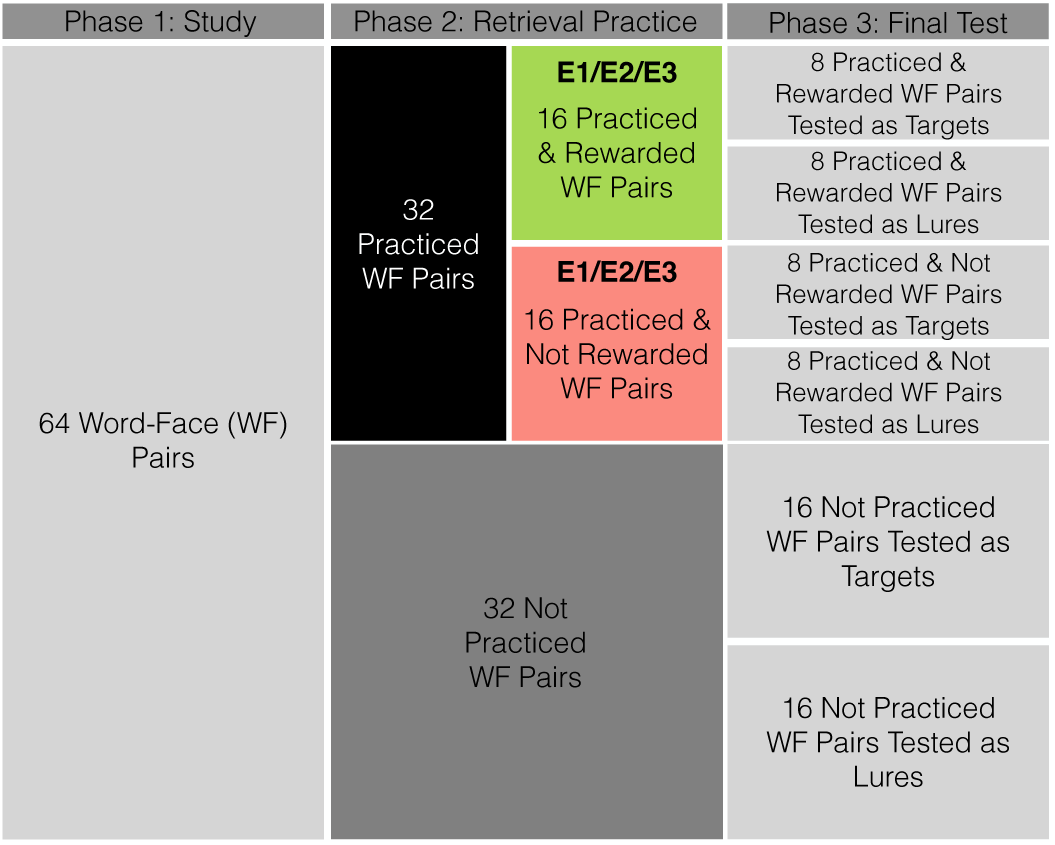
Condition Divisions and Counts. For each image category, 64 word-image pairs are shown during Phase 1 study. We show the specific example of word-face (WF) pairs here, but the same applies to word-object and word-scene pairs. Across all four studies, of the 64 studied WF pairs, 32 are practiced and 32 are not practiced. For E1, E2, and E3, the 32 practiced pairs are further sub-divided into 16 rewarded and 16 not rewarded practiced pairs. During the Phase 3 final test, pairs are tested as either targets (the exact image presented during study is presented during test) or lures (non-identical images depicting the same face as the Phase 1 image). Therefore, across all four experiments there are 16 not practiced pairs tested as targets and 16 not practiced pairs tested as lures. In E4 which has no rewards there are likewise 16 practiced pairs tested as targets/lures. In E1, E2, E3, there is further sub-division on the basis of reward yielding 8 pairs per bin (rewarded/not rewarded and tested as target/lure).

*Phase 1: Study.* In each of four runs, participants studied 48 word-image pairs, yielding a total of 192 trials. On each trial, participants saw a word-image pair presented for 2000 ms followed by a 3500 ms distractor interval. The distractor interval was comprised of alternating fixation and digit presentation (fixation, digit, fixation, digit, fixation). During digit presentation, participants saw a single digit, 1 through 10, and were instructed to press one of two buttons (“1” or “2”) to indicate if the number was odd or even. Each fixation was 500 ms and each digit presentation was 1000 ms.

*Phase 2: Retrieval Practice.* Participants completed two rounds of retrieval practice containing 96 trials each. A total of 96 words from Phase 1 were presented. The same words were presented in both rounds of retrieval practice in random order. Each round was further sub-divided into two runs with 48 trials.

In each run, an equal number of words (16) associated with an image from each visual category were presented. On each trial, participants were presented with a word cue for 4000 ms and instructed to bring to mind the associated image from the study phase. Participants rated the vividness of their memory of the retrieved image on a scale from 1 to 4, with 1 being least vivid and 4 being most vivid. To motivate participants to use the full response scale, if the same vividness response was made on five or more consecutive trials, participants received a message instructing them to use the full scale. Following the word cue, participants saw either a scrambled mask or feedback displayed as coins overlaid on the scrambled mask for 1000 ms, followed by a 500 ms inter-stimulus interval (ISI).

We manipulated reward delivery across experiments. In E1, participants received rewards during only the first round of retrieval practice. Every response had a 50% chance of receiving reward feedback. In E2, participants received rewards during only the second round of retrieval practice. As in E1, every response had a 50% chance of receiving reward feedback. In E3, participants received rewards during both rounds of retrieval practice. In the first round, as in E1, every response had a 50% chance of receiving reward feedback. If reward followed an item in the first round of retrieval practice, reward followed the same item in the second round. In E1, E2, and E3, 48 total words were rewarded (16 words associated with images from each visual category). In E4, participants did not receive any reward during either round of retrieval practice.

*Phase 3: Final Test.* Participants completed a final recognition memory test for images only. Trials were self-paced and participants made old or new judgements for each image using a confidence rating scale from 1 to 4, with 1 being definitely new and 4 being definitely old. Trials were separated by a 500 ms ISI. There were a total of 288 test trials. Test probes included 96 previously studied images (targets), 96 highly similar lures (non-identical images depicting the same face, scene, or object as the Phase 1 image), and 96 novel face, scene, or object images. To reduce test phase interference, participants were only tested on either a target (e.g. original image of Britney Spears) or the similar lure (e.g. new image of Britney Spears), as in prior work (Lee et al., 2018). There were an equal number of images from each visual category for each test probe condition (i.e. 32 novel scenes). Half (48) of the targets were practiced and half were not practiced; likewise, half of the lures were associated with an image that was practiced and half were associated with an image that was not practiced. We refer to these as “practiced lures” and “not practiced lures” although the lures themselves were not practiced (Figure 1B). For E1, E2, and E3, half (24) of the practiced targets were rewarded during retrieval practice and half were not rewarded. Similarly, half of the practiced lures were associated with an image that was rewarded during retrieval practice and half were associated with an image that was not rewarded during retrieval practice. We refer to these as “rewarded targets” and “rewarded lures” although the rewards were always presented during the Phase 2 retrieval practice and never during the final Phase 3 recognition test.

### Statistical Analyses

We used mixed effects ANOVAs to assess the effect of retrieval practice, reward structure, and vividness on hit rate and false alarm rate. We used an independent samples *t* -test to compare E1-E3 (rewarded and not rewarded) hit rate and false alarm rate to E4 hit rate and false alarm rate.

## Results

### Reward structure impacts subsequent hit rates

Our first goal was to test whether rewarded practice, regardless of the precise structure of reward delivery, modulates memory performance. Following our pre-registration, we collapsed data from the three experiments with reward (E1, E2, E3) and compared hit and false alarm (FA) rates from these experiments to hit and FA rates in the control experiment (E4) that had no rewards (Figure 3). Insofar as extrinsic rewards reinforce the contents of retrieval, we expected to find higher hit rates and lower FA rates for rewarded compared to control trials. We used independent sample *t* -tests to compare hit rates and false alarm rates and find no significant differences across the rewarded experiments and the control experiment (Table 1). Thus, we do not find evidence that rewards during retrieval practice impact recognition memory performance.

**Figure 3.**
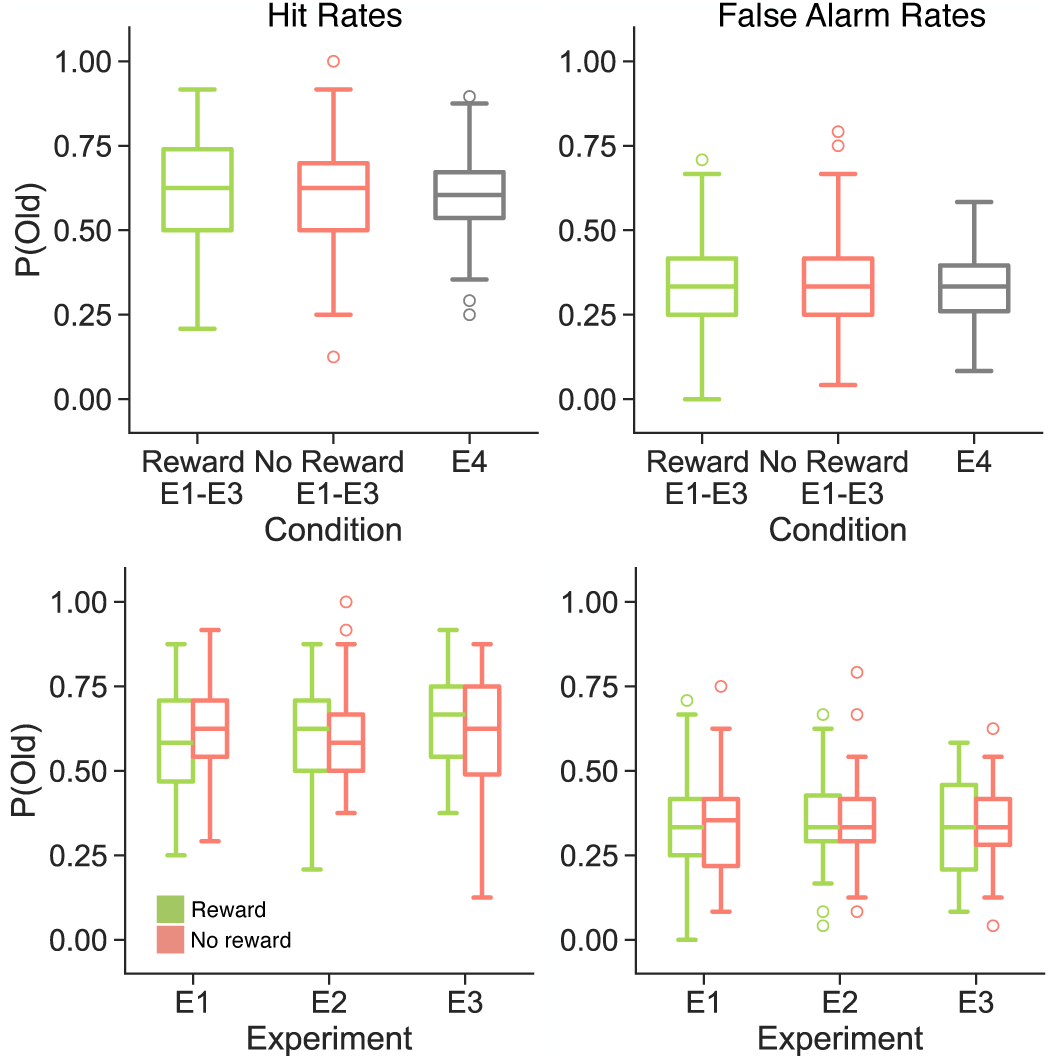
Influence of reward delivery and reward structure on hit and false alarm rates. Each panel shows the proportion of old responses to reward (green), no reward (red) and/or E4 (grey) probes. The left columns show hit rates and the right columns show false alarm rates. The top row shows data averaged over experiments 1-3; the bottom row shows data separately for each experiment. Box-and-whisker plots show median (center line), upper and lower quartiles (box limits), 1.5x interquartile range (whiskers) and outliers (circles). We do not find a significant difference in either hit rates or false alarm rates for reward or no reward E1-E3 probes compared to E4 probes. We find a significant interaction between reward and experiment (p = 0.0385) on hit rates, but not on false alarm rates (p = 0.9505).

**Table 1.** Memory performance for rewarded/not rewarded practice (E1/E2/E3) compared to control (E4).

|  | Hits |  | vs. Control |  |  |  | FAs |  | vs. Control |  |  |  |
| --- | --- | --- | --- | --- | --- | --- | --- | --- | --- | --- | --- | --- |
| | Mean | SD | $t_{136}$ | $p$ | $d$ | $BF$ | Mean | SD | $t_{136}$ | $p$ | $d$ | $BF$ |
| Rewarded | 0.6132 | 0.1561 | 0.536 | 0.5929 | 0.1072 | 0.2331 | 0.3411 | 0.1428 | 0.2046 | 0.8382 | 0.0418 | 0.2089 |
| Not rewarded | 0.5993 | 0.1583 | 0.068 | 0.9459 | 0.0136 | 0.2055 | 0.3452 | 0.141 | 0.3616 | 0.7182 | 0.0738 | 0.2174 |
| Control | 0.5972 | 0.1408 | - | - | - | - | 0.3356 | 0.1162 | - | - | - | - |

Despite failing to find a general difference between rewarded practice and the control experiment, we expected that reward structure might differentially impact subsequent memory. Specifically, we expected that repeated rewards (E3) might increase subsequent hit rates relative to singly presented rewards (E1, E2). Additionally, we expected that lack of rewards in the second round of practice (E1) might decrease subsequent hit rates. To directly test the effect of reward structure on subsequent hit rates, we performed a 2 *×* 3 mixed effects ANOVA with reward delivery (rewarded, not rewarded) and experiment (E1, E2, E3) as factors, following our pre-registration (Figure 3). We do not find a significant main effect of either reward delivery (*F* _1,99_ = 0.92, *p* = 0.341, *η_p_*^2^ = 0.009) or experiment (*F* _2,99_ = 0.16, *p* = 0.852, *η_p_*^2^ = 0.003). We find a significant reward delivery by experiment interaction (*F* _1,99_ = 3.37, *p* = 0.039, *η_p_*^2^ = 0.064). These results suggest that the structure of rewards – how many and when they are received – impacts subsequent memory.

As we anticipated a difference between rewards presented once vs. twice, we conducted two follow-up ANOVAs in which we compared E1 and E2 to E3 (Table 2). For the E1 vs. E3 ANOVA, we find a significant reward by experiment interaction, driven by significantly greater hit rates for rewarded vs. not rewarded targets in E3 (rewarded, M = 0.6510, SD = 0.1408; not rewarded, M = 0.5846, SD = 0.1803, *t* _31_ = 2.51, *p* = 0.018, *d* = 0.411, BF = 2.755) and numerically lower hit rates for rewarded vs. not rewarded targets in E1 (rewarded, M = 0.5882, SD = 0.1613; not rewarded, M = 0.6140, SD = 0.1567, *t* _33_ = -0.92, *p* = 0.366, *d* = 0.162, BF = 0.271). For the E2 vs. E3 ANOVA, we find a main effect of reward, whereby hit rates are greater for rewarded compared to not rewarded trials. Although numerically hit rates were greater for rewarded vs. not rewarded targets in E2 (rewarded, M = 0.6030, SD = 0.1576; not rewarded, M = 0.5984, SD = 0.1361), this difference was not significant (*t* _35_ = 0.22, *p* = 0.826, *d* = 0.031, BF = 0.183).

**Table 2.** Hit rates as a function of reward structure.

| Effect | E1 vs. E3 |  |  |  | E2 vs. E3 |  |  |  |
| --- | --- | --- | --- | --- | --- | --- | --- | --- |
| | df | $F$ | $p$ | $\eta_p^2$ | df | $F$ | $p$ | $\eta_p^2$ |
| Main effect of reward | (1,64) | 0.959 | 0.3310 | 0.01 | (1,66) | **4.105** | **0.0468** | **0.06** |
| Main effect of experiment | (1,64) | 0.227 | 0.635 | 0.003 | (1,66) | 0.251 | 0.618 | 0.003 |
| Interaction of reward × experiment | (1,64) | **5.671** | **0.0202** | **0.08** | (1,66) | 3.436 | 0.0682 | 0.05 |

*Note: Bold values indicate $p < 0.05$*

Following our pre-registration, we performed the same 2 *×* 3 mixed effects ANOVA on FA rates (Figure 3). We do not find a significant main effect of reward delivery (*F* _1,99_ = 0.11, *p* = 0.741, *η_p_*^2^=0.001) or experiment (*F* _2,99_ = 0.33, *p* = 0.723, *η_p_*^2^= 0.007). We do not find a significant reward delivery by experiment interaction (*F* _1,99_ = 0.05, *p* = 0.951, *η_p_*^2^= 0.001).

Taken together, these findings suggest that repeated reward during retrieval practice can facilitate later target detection.

### Vividness during retrieval practice differentially impacts subsequent memory

Given the relatively limited impact of rewarded retrieval practice on subsequent memory, we next conducted a series of exploratory analyses in which we investigated the effect of practice and vividness on subsequent memory. The modest effect of rewarded practice on subsequent memory may due to strong effects of either practice itself and/or the vividness experienced during practice.

We first tested the extent to which retrieval practice modulates hit and FA rates (Figure 4). Based on extensive prior work (e.g. Roediger & Karpicke, 2006), we expected to find higher hit and FA rates for practiced relative to not practiced items. We conducted two paired-samples *t* -tests on data aggregated across all four studies and do not find a significant effect of practice for either hit rates (practiced, M = 0.6039, SD = 0.1390; not practiced, M = 0.6004, SD = 0.1425; *t* _137_ = 0.45, *p* = 0.654, *d* = 0.025, BF = 0.105) or FA rates (practiced, M = 0.3412, SD = 0.1250; not practiced, M = 0.3474, SD = 0.1186; *t* _137_ = -0.84, *p* = 0.402, *d* = 0.051, BF = 0.134). Thus we do not find evidence that practice alone modulates subsequent memory.

**Figure 4.**
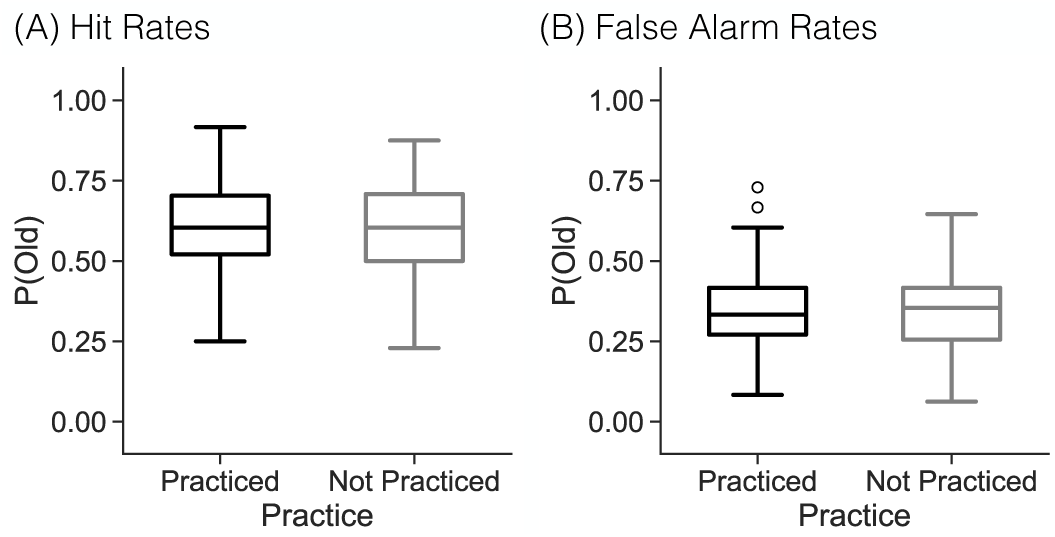
Influence of retrieval practice on hit and false alarm rates. Each panel shows the proportion of old responses to practice (black) and not practiced (grey) probes. Box-and-whisker plots show median (center line), upper and lower quartiles (box limits), 1.5x interquartile range (whiskers) and outliers (circles). **(A)** We do not find a significant difference in hit rates between practice and not practiced targets (p = 0.6535). **(B)** We do not find a significant difference in false alarm rates between practice and not practiced lures (p = 0.4015).

We were somewhat surprised that we did not find a significant effect of retrieval practice on subsequent memory. However, we speculate that the vividness with which participants remembered the associated image stimuli may influence the effect of retrieval practice on subsequent memory. In particular, given prior evidence that partial or incomplete reinstatement (Newman & Norman, 2010; Kim, Lewis-Peacock, Norman, & Turk-Browne, 2014; Kim, Norman, & Turk-Browne, 2017) can negatively impact subsequent memory, there may be a bidirectional effect of practice: enhancement for high vivid items and decrement for low vivid items which when aggregated would produce the observed null effect of retrieval practice. To test this possibility, we compared hit rates and FA rates across three conditions: practiced items that received a high vividness rating (3 or 4, “high vivid”), practiced items that received a low vividness rating (1 or 2, “low vivid”), and not practiced items (Figure 5). We specifically used vividness ratings from the first round of retrieval practice across all experiments as these initial ratings could not be modulated by reward or practice itself. We conducted two 1 *×* 3 rmANOVA with three levels: high vivid, low vivid, and not practiced. We expected to find that hit rates and FA rates would increase for high vivid compared to not practiced items, but would decrease for low vivid compared to not practiced items. We find a significant effect of condition for both hits and FAs (hits: *F* _2,274_ = 127.10, *p <* 0.0001, *η_p_*^2^ = 0.481; FAs: *F* _2,272_ = 25.79, *p <* 0.0001, *η_p_*^2^ = 0.159, *Note: one participant was excluded from the FA analysis for not having any high vivid FAs*). In both cases, these effects were driven by significantly greater hit rates and FA rates for high vivid compared to not practiced items and for not practiced items compared to low vivid items (Table 3).

**Figure 5.**
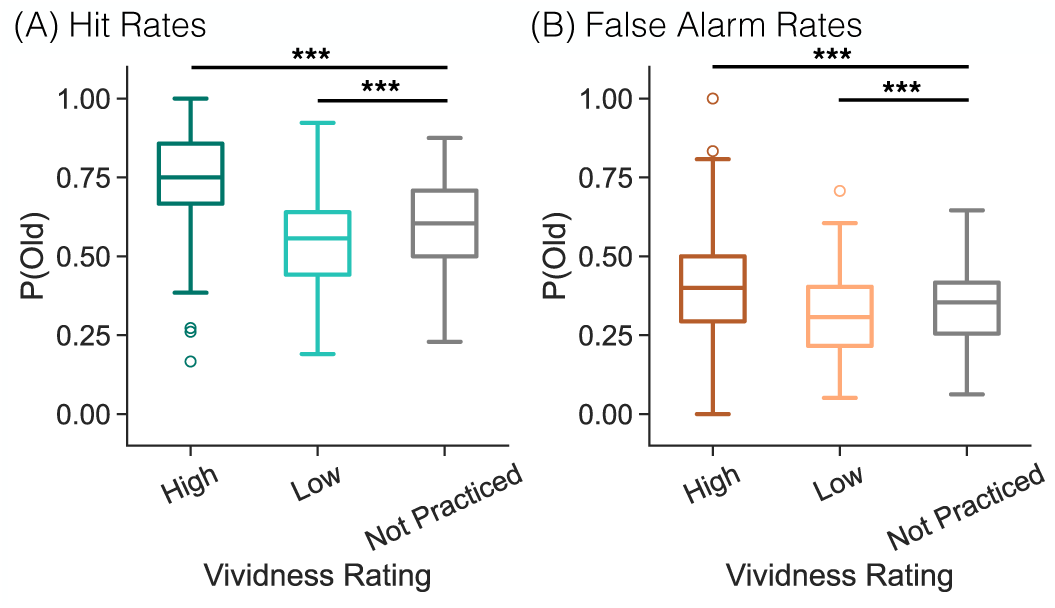
Influence of vividness on hit and false alarm rates. Each panel shows the proportion of old responses to high vivid (ratings of 3 or 4, darker), low vivid (ratings of 1 or 2, lighter), and not practiced (grey) probes. Box-and-whisker plots show median (center line), upper and lower quartiles (box limits), 1.5x interquartile range (whiskers) and outliers (circles). *** p *<* 0.0001 **(A)** We find a significant effect of vividness driven by greater hit rates for high vivid (dark teal) compared to not practiced targets and for not practiced compared to the low vivid (light teal) targets. (**B**) We find a significant main effect of vividness driven by greater false alarm rates for high vivid (dark orange) compared to not practiced lures and for not practiced compared to the low vivid (light orange) lures.

**Table 3.** Memory performance as a function of practice vividness.

|  | Hits |  | vs. Not Practiced |  |  |  | FAs |  | vs. Not Practiced |  |  |  |
| --- | --- | --- | --- | --- | --- | --- | --- | --- | --- | --- | --- | --- |
| | Mean | SD | $t_{137}$ | $p$ | $d$ | $BF$ | Mean | SD | $t_{136}$ | $p$ | $d$ | $BF$ |
| High Vivid | 0.7349 | 0.1785 | **10.59** | **<0.0001** | **0.8328** | $1.0 \times 10^{18}$ | 0.4177 | 0.1877 | **4.741** | **<0.0001** | **0.4411** | 300.5 |
| Low Vivid | 0.5532 | 0.1507 | **-4.608** | **<0.0001** | **0.3216** | $4.7 \times 10^4$ | 0.3156 | 0.1364 | **-3.819** | **0.0002** | **0.2583** | 77.11 |
| Not Practiced | 0.6004 | 0.1425 | - | - | - | - | 0.3485 | 0.1182 | - | - | - | - |

*Note: Bold values indicate $p < 0.05$*

### Rewards interact with vividness to modulate subsequent memory

As we have separately shown that repeated rewards (E3) and vividness during practice influence later memory, our final goal was to evaluate the joint impact of rewards and vividness. We focus specifically on data from the first round of practice in E3, given that vividness ratings from this round of practice are unaffected by reward and practice. Furthermore, we focus specifically on hit rates given that the structure of reward delivery had a significant effect on hit rates. We separately consider subsequent hit rates for high and low vivid items vs. not practiced items. To the extent that vividness judgments reflect the degree of reinstatement for a given representation (e.g. Kuhl & Chun, 2014), providing a reward following a highly vivid retrieval might have little impact on the associated neural representation which is presumably already strong or of high fidelity. However, to the extent that a low vivid response during retrieval practice reflects a weak or low fidelity representation, reward may impact said representation and thus later memory.

We performed two 1 *×* 3 ANOVAs to separately compare high vivid (rewarded, not rewarded) to not practiced items and low vivid (rewarded, not rewarded) to not practiced items (Figure 6) and found a significant effect for each (high vivid: *F* _2,62_ = 11.81, *p <* 0.0001, *η_p_*^2^ = 0.28; low vivid: *F* _2,62_ = 5.50, *p* = 0.0006, *η_p_*^2^ = 0.15). We conducted post-hoc paired *t* -tests separately for the high and low vivid comparisons. We find that hit rates are significantly greater for both high vivid rewarded (M = 0.8173, SD = 0.1614) and high vivid not rewarded (M = 0.6977, SD = 0.2846) items compared to not practiced items (rewarded: *t* _31_ = 7.78, *p <* 0.0001, *d* = 1.28, BF = 1.40 *×* 10^6^; not rewarded: *t* _31_ = 2.13, *p* = 0.042, *d* = 0.362, BF = 1.352). We find that hit rates for low vivid rewarded items (M = 0.6016, SD = 0.1693) do not significantly differ from hit rates for not practiced items (M = 0.6146, SD = 0.1566, *t* _31_ = -0.48, *p* = 0.632, *d* = 0.080, BF = 0.211). We find that hit rates for low vivid not rewarded items (M = 0.5346, SD = 0.1672) are significantly lower than hit rates for not practiced items (*t* _31_ = -4.37, *p* = 0.0001, *d* = 0.494, BF = 200). Together, these results suggest that rewards may ‘recover’ or amplify items that are only partially reinstated or remembered with low vividness.

**Figure 6.**
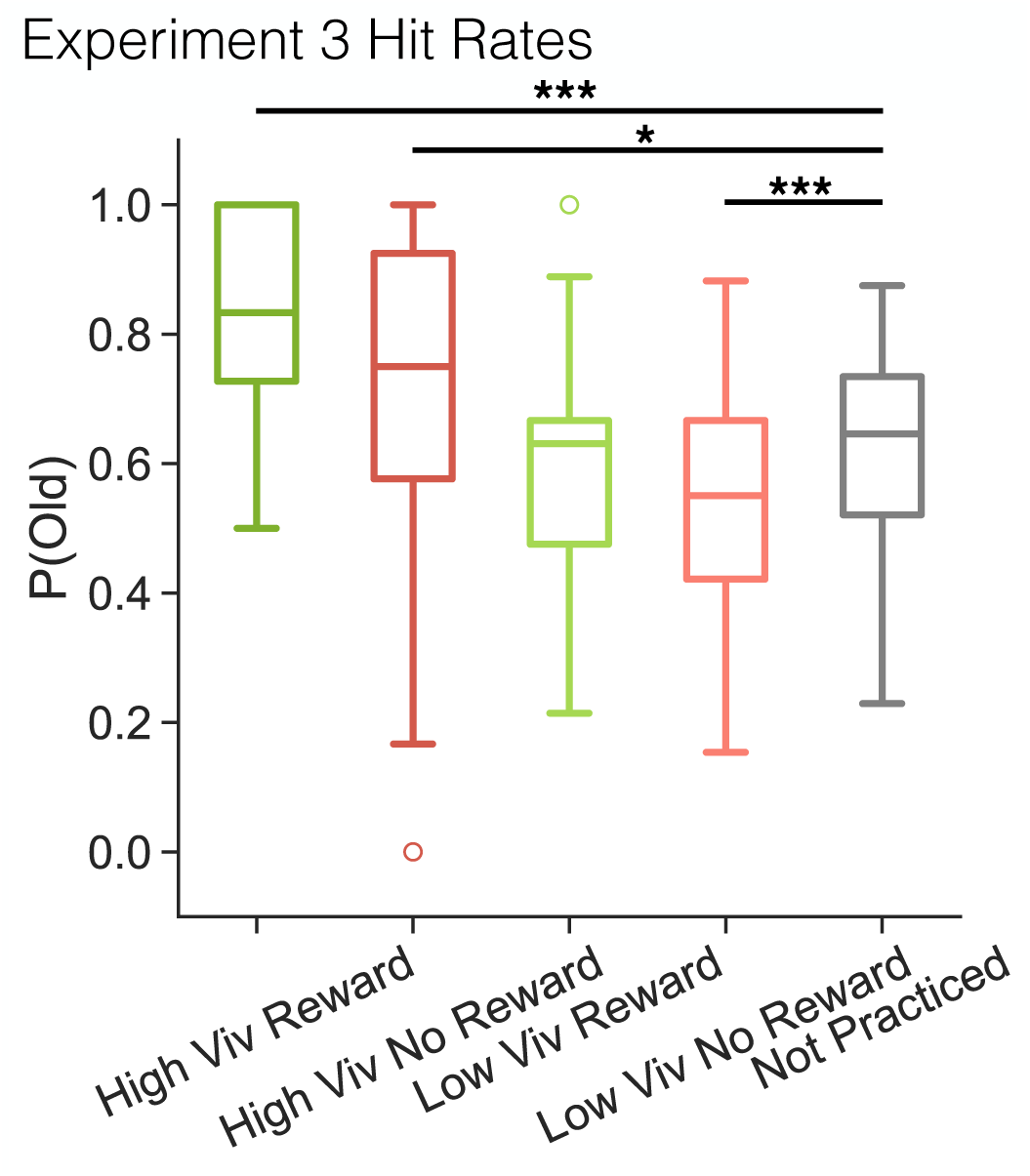
Influence of vividness, reward, and practice on hit rates in E3. We find a significant difference in hit rates between low vivid targets that were not rewarded and not practiced targets. Box-and-whisker plots show median (center line), upper and lower quartiles (box limits), 1.5x interquartile range (whiskers) and outliers (circles). * p *<* 0.05, *** p *<* 0.0001

## Discussion

The aim of this study was to investigate how extrinsic reward following retrieval practice impacts subsequent memory. We conducted four independent recognition memory experiments in which we measured the impact of reward delivery, reward structure, retrieval practice, and vividness on recognition memory. We find that repeated rewards increase subsequent hit rates relative to singly presented rewards and potentially amplify low fidelity memory representations. Together, these findings suggest that extrinsic rewards following retrieval have the potential to benefit later memory.

We find that the structure of reward delivery during retrieval practice modulates subsequent memory. Specifically, we find higher hit rates for practice items that were rewarded during both of two rounds of retrieval practice. As rewards in the current study were random, it is unlikely that participants engaged particular strategies prior to specific word cues as in traditional anticipatory designs (e.g. Castel, 2007).

Instead, we speculate that random rewards acted on the reactivated representations of the associated images. Individuals reactivate both item-specific and category level information while making vividness judgments (e.g. Kuhl et al., 2013) and surprising rewards can promote enhanced connectivity between task-specific regions, the hippocampus, and the striatum (Calderon et al., 2021). Potentially, each reward presentation may offer the opportunity to strengthen reactivated representations, leading to the memory facilitation that we find for twice-presented rewards. Although future neuro-imaging work is needed to directly test the link between cortical reactivation and reward system activation, our findings are consistent with the interpretation that post-retrieval rewards may act directly on reactivated memories. Alternatively, rewards may induce a content-general increase in attention to the lingering internal representation. Sustained attention may extend the window of what is effectively test-phase encoding (Nyberg, Habib, & Tulving, 2000), facilitating later memory.

Although we do not find a main effect of retrieval practice, we find robust effects of retrieval practice vividness on subsequent memory. We replicate prior work and find that high vividness ratings during retrieval practice are associated with greater hit rates and false alarm rates relative to not practiced items (Lee et al., 2018). We also find lower hit rates for items low in vividness. Our interpretation is that the consequences of retrieval practice vary depending on the strength of the information that is retrieved. Our findings are in line with the non-monotonic plasticity hypothesis (Poppenk & Norman, 2014), whereby moderate reactivation of an item may lead to the representation of that item being weakened (Newman & Norman, 2010; Kim et al., 2014). Thus, it may be that low vivid items in the present study were partially reactivated and therefore weakened by practice.

Repeated post-retrieval rewards improve subsequent memory for low vivid items. We find that practiced items that receive a low vividness rating have lower subsequent hit rates compared to items that are not practiced at all. However, we find that subsequent memory is comparable for low vivid practice *rewarded* items relative to not practiced items. Thus, whereas low vivid items may be weakened through retrieval practice (Poppenk & Norman, 2014), rewards may counteract this weakening, potentially by amplifying below-threshold signals – moving the reactivated memory out of the ‘moderate’ zone and into a higher reactivation state.

Our use of random rewards was motivated by several factors; however, random rewards are not strongly ecologically plausible. We used random rewards so as to not link rewards to specific behaviors (e.g. correct memory of the associated image), contexts (e.g. a particular image category), and/or motor responses (e.g. a button press of 4). Such a structure would likely motivate participants to respond ‘4’ to all retrieval practice trials and/or only attend to a specific category (e.g. faces). Additionally, and perhaps most critically, our goal was to *not* induce pre-retrieval reward anticipation. Reward anticipation can influence participant strategies and decisions (Castel, 2007; Bowen, Marchesi, & Kensinger, 2020) such that non-random rewards would likely alter participants’ processing of the word cues rather than, or in addition to, processing of the reinstated image. Such alterations will have differential impacts on later memory, especially depending on the format of the memory test. In the current study, we used a recognition test to assess memory for images whose representations were theoretically modulated by rewarded retrieval practice. In contrast, an anticipatory design may serve to facilitate word-image associations instead. However, truly random rewards are unlikely in the real world where rewards are instead linked to specific behaviors. Reward prediction errors (RPEs) – when the predicted outcome deviates from what is received – are well known to impact behavior (Schultz, Dayan, & Montague, 1997). As both positive and negative RPEs – receiving unexpected rewards or unexpected punishments – modulate neural signals and drive behavior (Zaghloul et al., 2009; Scimeca, Katzman, & Badre, 2016; Jang, Nassar, Dillon, & Frank, 2019; Ergo, De Loof, & Verguts, 2020; Rouhani & Niv, 2021), RPEs during retrieval practice may reinforce memories through different mechanisms than random rewards. Future work will be needed to directly test this possibility.

Extrinsic rewards can negatively impact behavior by altering arousal levels (Cheng et al., 2020) and interfering with intrinsic rewards (Deci, Koestner, & Ryan, 1999). Although we cannot draw strong conclusions on the basis of null results, given the lack of an effect of rewarded retrieval practice on false alarms, we speculate that extrinsic rewards in the current study are likely reinforcing both item and category reactivation. In contrast to item-specific reactivation, category reactivation promotes false alarms (Lee et al., 2018). Thus, post-retrieval rewards could be both beneficial and maladaptive depending on the exact contents of retrieval. Future neuro-imaging work will be needed to address this possibility. More broadly, extrinsic rewards can diminish the impact of internal motivation (Dickerson & Adcock, 2018; Xue et al., 2023). To the extent that successful retrieval is intrinsically rewarding (Speer et al., 2014; Smith & Long, 2024), extrinsic rewards during retrieval practice may interfere with intrinsic reward signals and inconsistently impact later memory.

Taken together, our findings demonstrate that rewarded retrieval practice facilitates subsequent recognition. In particular, low vivid retrieval practice items – which would typically be remembered worse than not practiced items – appear to be ‘recovered’ by random extrinsic rewards. These findings suggest that the reward system may interact with, and reinforce, the contents of retrieval, impacting subsequent memory. More broadly, we contribute to a growing body of literature showing a strong interconnection between memory and reinforcement learning.

## Research Transparency Statement

### General Disclosures

Conflicts of interest: None. Funding: None. Artificial intelligence: None. Ethics: This research received approval from the University of Virginia Social Behavioral Sciences Institutional Review Board (protocol #2115).

Preregistration: the hypotheses, methods, and analysis plan were preregistered (https://osf.io/gebm4) prior to data collection. All pre-registered analyses were performed. Additional exploratory analyses, not preregistered, were also performed. Materials: All study materials are publicly available (osf.io/y5ac8). Data: All primary data are publicly available (osf.io/y5ac8). Analysis scripts: All analysis scripts are publicly available (osf.io/y5ac8).

## CRediT authorship contribution statement

**Devyn E. Smith:** Conceptualization, Data Curation, Formal analysis, Investigation, Methodology, Software, Validation, Visualization, Writing – original draft, Writing – review & editing. **Amanda M. Smith:** Formal analysis, Investigation, Software, Validation, Visualization, Writing – review & editing. **Hannah R. Buras:** Data Curation, Investigation, Writing – review & editing. **Nicole M. Long:** Conceptualization, Formal analysis, Funding acquisition, Methodology, Project administration, Resources, Software, Supervision, Validation, Visualization, Writing – review & editing.

## References

1. Adcock, A. R., Thangavel, A., Whitfield-Gabrieli, S., Knutson, B., & Gabrieli, J. D. (2006). Reward-motivated learning: Mesolimbic activation precedes memory formation. Neuron, 50, 507–517.

2. Bowen, H. J., Marchesi, M. L., & Kensinger, E. A. (2020). Reward motivation influences response bias on a recognition memory task. Cognition, 203, 1–13.

3. Brainerd, C., & Reyna, V. (2002). Fuzzy-trace theory and false memory. Current Directions in Psychological Science, 11(5), 164–169.

4. Calderon, C. B., De Loof, E., Ergo, K., Snoeck, A., Boehler, C. N., & Verguts, T. (2021). Signed reward prediction errors in the ventral striatum drive episodic memory. Journal of Neuroscience, 41(8), 1716–1726.

5. Castanheira, K. d. S., Lalla, A., Ocampo, K., Otto, A. R., & Sheldon, S. (2022). Reward at encoding but not retrieval modulates memory for detailed events. Cognition, 219(104957), 1–10.

6. Castel, A. D. (2007). The adaptive and strategic use of memory by older adults: Evaluative processing and value-directed remembering. In Psychology of learning and motivation (Vol. 48). Elsevier.

7. Castel, A. D., Benjamin, A. S., Craik, F. I., & Watkins, M. J. (2002). The effects of aging on selectivity and control in short-term recall. Memory & Cognition, 30(7), 1078–1085.

8. Cheng, S., Jiang, T., Xue, J., Wang, S., Chen, C., & Zhang, M. (2020). The influence of rewards on incidental memory: more does not mean better. Learning & Memory . doi: 10.1101/lm.051722.120

9. Chung, Y. M. W., & Federmeier, K. D. (2023). Read carefully, because this is important! how value-driven strategies impact sentence memory. Memory & Cognition, 51(7).

10. Deci, E. L., Koestner, R., & Ryan, R. M. (1999). A meta-analytic review of experiments examining the effects of extrinsic rewards on intrinsic motivation. Psychological Bulletin, 125(6), 627–668.

11. Detre, G. J., Natarajan, A., Gershman, S. J., & Norman, K. A. (2013). Moderate levels of activation lead to forgetting in the think/no-think paradigm. Neuropsychologia, 51(12), 2371–2388.

12. Diana, R. A., Yonelinas, A. P., & Ranganath, C. (2007). Imaging recollection and familiarity in the medial temporal lobe: a three-component model. TRENDS in Cognitive Sciences, 11(9), 379–386.

13. Dickerson, K. C., & Adcock, R. A. (2018). Motivation and memory. In J. T. Wixted (Ed.), Steven’s handbook of experimental psycholgoy and cognitive neuroscience. John Wiley & Sons, Inc.

14. Eichenbaum, H. (2004). Hippocampus: Cognitive processes and neural representations that underlie declarative memory. Neuron, 44, 109–120.

15. Elliott, B. L., Blais, C., McClure, S. M., & Brewer, G. A. (2020). Neural correlates underlying the effect of reward value on recognition memory. NeuroImage, 206, 1–9.

16. Ergo, K., De Loof, E., & Verguts, T. (2020). Reward prediction error and declarative memory. Trends in Cognitive Sciences, 24(5), 388–397.

17. Filiz, G., & Dobbins, I. G. (2024). The limited memory of value following value directed encoding. Memory & Cognition. doi: 10.3758/s13421-024-01550-7

18. Friendly, M., Franklin, P. E., Hoffman, D., & Rubin, D. C. (1982). The toronto word pool: Norms for imagery, concreteness, orthographic variables, and grammatical usage for 1,080 words. Behavior Research Methods & Instrumentation, 14, 375–399.

19. Jang, A. I., Nassar, M. R., Dillon, D. G., & Frank, M. J. (2019). Positive reward prediction errors during decision making strengthen memory encoding. Nature human behavior, 3(7), 719–732.

20. Karpicke, J. D. (2012). Retrieval-based learning: Active retrieval promotes meaningful learning. Current Directions in Psychological Science, 21(3), 157–163.

21. Karpicke, J. D., & Roediger, H. L. (2008). The critical importance of retrieval for learning. Science, 319, 966–968.

22. Kim, G., Lewis-Peacock, J. A., Norman, K. A., & Turk-Browne, N. B. (2014). Pruning of memories by context-based prediction error. Proceedings of the National Academy of Sciences of the United States of America, 111(24).

23. Kim, G., Norman, K. A., & Turk-Browne, N. B. (2017). Neural differentiation of incorrectly predicted memories. Journal of Neuroscience, 37 (8), 2022–2031.

24. Knowlton, B. J., & Castel, A. D. (2022). Memory and reward-based learning: A value-directed remembering perspective. Annual Review of Psychology, 73(1), 25–52.

25. Konkle, T., Brady, T. F., Alvarez, G. A., & Oliva, A. (2010). Conceptual distinctiveness supports detailed visual long-term memory for real-world objects. Journal of Experimental Psychology: General, 139(3), 558–578.

26. Kuhl, B. A., & Chun, M. M. (2014). Successful remembering elicits event-specific activity patterns in lateral parietal cortex. The Journal Of Neuroscience, 34(23), 8051–8060.

27. Kuhl, B. A., Johnson, M. K., & Chun, M. M. (2013). Dissociable neural mechanisms for goal-directed versus incidental memory reactivation. The Journal of Neuroscience, 33(41), 16099–16109.

28. Lee, H., Samide, R., Richter, F. R., & Kuhl, B. A. (2018). Decomposing parietal memory reactivation to predict consequences of remembering. Cerebral Cortex, 29, 1–14.

29. Loftus, G. R., & Wickens, T. D. (1970). Effect of incentive on storage and retrieval processes. Journal of Experimental Psychology, 85(1), 141–147.

30. Long, N. M., Sperling, M. R., Worrell, G. A., Davis, K. A., Gross, R. E., Lega, B. C., . . . Kahana, M. J. (2017). Contextually mediated spontaneous retrieval is specific to the hippocampus. Current Biology, 27, 1–6.

31. Marini, F., Marzi, T., & Viggiano, M. P. (2011). “wanted!” the effects of reward on face recognition: electrophysiological correlates. Cogn Affect Behav Neurosci, 11, 627–643.

32. McDermott, K. B. (2006). Paradoxical effects of testing: repeated retrieval attempts enhance the likelihood of later accurate and false recall. Memory & Cognition, 34(2), 261–267.

33. Newman, E. L., & Norman, K. A. (2010). Moderate excitation leads to weakening of perceptual representations. Cerebral Cortex, 20(11), 2760–2770.

34. Nyberg, L., Habib, R., & Tulving, E. (2000). Reactivation of encoding-related brain activity during memory retrieval. Proceedings of the National Academy of Sciences of the United States of America, 97, 11120.

35. Oyarzún, J. P., Packard, P. A., de Diego-Balaguer, R., & Fuentemilla, L. (2016). Motivated encoding selectively promotes memory for future inconsequential semantically-related events. Neurobiology of learning and memory, 133, 1–6.

36. Poppenk, J., & Norman, K. A. (2014). Briefly cuing memories leads to suppression of their neural representations. The Journal of Neuroscience, 34(23), 8010–8020.

37. Roediger, H. L., & Karpicke, J. D. (2006). Test-enhanced learning: taking memory tests improves longterm retention. Psychological Science, 17 (3), 249–255.

38. Rouhani, N., & Niv, Y. (2021). Signed and unsigned reward prediction errors dynamically enhance learning and memory. eLife, 10(e61077), 1–28.

39. Schultz, W., Dayan, P., & Montague, P. R. (1997). A neural substrate of prediction and reward. Science, 275(5306), 1593–1599.

40. Scimeca, J. M., Katzman, P. L., & Badre, D. (2016). Striatal prediction errors support dynamic control of declarative memory decisions. Nature Communications, 7 (13061), 1–15.

41. Shigemune, Y., Tsukiura, T., Nouchi, R., Kambara, T., & Kawashima, R. (2017). Neural mechanisms underlying the reward-related enhancement of motivation when remembering episodic memories with high difficulty. Human Brain Mapping, 38, 3428–3443.

42. Smith, D. E., & Long, N. M. (2024). Successful retrieval is its own reward. bioRxiv . doi: 10.1101/2024.07.26.605274

43. Speer, M. E., Bhanji, J. P., & Delgado, M. R. (2014). Savoring the past: positive memories evoke value representations in the striatum. Neuron, 84(4), 847–856.

44. Wimmer, G. E., & Shohamy, D. (2012). Preference by association: How memory mechanisms in the hippocampus bias decisions. Science, 338.

45. Wolosin, S. M., Zeithamova, D., & Preston, A. R. (2012). Reward modulation of hippocampal subfield activation during successful associative encoding and retrieval. Journal of Cognitive Neuroscience, 24(7), 1532–1547.

46. Xue, J., Jiang, T., Chen, C., Murty, V. P., Li, Y., Ding, Z., & Zhang, M. (2023). The interactive effect of external rewards and self-determined choice on memory. Psychological Research.

47. Zaghloul, K. A., Blanco, J. A., Weidemann, C. T., McGill, K., Jaggi, J. L., Baltuch, G. H., & Kahana, M. J. (2009). Human substantia nigra neurons encode unexpected financial rewards. Science, 323(5920), 1496–1499.

